# Micronized curcumin causes hyperlocomotion in zebrafish larvae

**DOI:** 10.1101/2021.11.29.470475

**Authors:** Adrieli Sachett, Radharani Benvenutti, Carlos G. Reis, Matheus Gallas-Lopes, Leonardo M. Bastos, Gean Pablo S. Aguiar, Ana P. Herrmann, J. Vladimir Oliveira, Anna M. Siebel, Angelo Piato

## Abstract

Zebrafish larvae have been widely used in neuroscience and drug research and development. In the larval stage, zebrafish present a broad behavioral repertoire and physiological responses similar to adults. Curcumin (CUR), a major component of *Curcuma longa* L. (Zingiberaceae), has demonstrated the ability to modulate several neurobiological processes relevant to mental disorders in animal models. However, the low bioavailability of this compound can compromise its *in vivo* biological potential. Interestingly, it has been shown that micronization can increase the biological effects of several compounds. Thus, in this study, we compared the effects of acute exposure for 30 minutes to the following solutions: water (control), 0.1% DMSO (vehicle), 1 μM CUR, or 1 μM micronized curcumin (MC) in zebrafish larvae 7 days post-fertilization (dpf). We analyzed locomotor activity (open tank test), anxiety (light/dark test), and avoidance behavior (aversive stimulus test). Moreover, we evaluated parameters of oxidative status (thiobarbituric acid reactive substances and non-protein thiols levels). MC increased the total distance traveled and absolute turn angle in the open tank test. There were no significant differences in the other behavioral or neurochemical outcomes. The increase in locomotion induced by MC may be associated with a stimulant effect on the central nervous system, which was evidenced by the micronization process.

## INTRODUCTION

Zebrafish have been used as a model organism for biomedical research in drug discovery, developmental biology, and genetics as it presents high mammalian homology [1]. Zebrafish present rapid development and a relatively long lifespan, being used to investigate the neurobehavioral foundations of various neurological and psychiatric conditions such as epilepsy, Alzheimer’s and Parkinson’s disease, schizophrenia, and stress- and drug-related disorders [1–9].

The low cost of rearing and maintenance and the numerous offspring are some of the attractive features of this organism when compared to rodents. This organism has proven to be very versatile in research, being used at different stages of development. Zebrafish larvae have a rich behavioral repertoire and respond to pharmacological and non-pharmacological interventions similarly to rodents [1, 8]. Studies have shown that zebrafish larvae respond to different classes of psychotropic drugs including ethanol, hallucinogens, and nicotine [10–15]. All these aspects support the use of zebrafish larvae in high-throughput screening for research and development of new drugs as well as for repurposing [2, 16].

Curcumin is the major active component extracted from the roots of turmeric *Curcuma longa* L. (Zingiberaceae). In larval and adult zebrafish, it has shown antioxidant [17–20], anti-seizure [5, 21], anti-inflammatory [20] and cognitive enhancing [22] activities. However, curcumin has low bioavailability *in vivo*, compromising its use [23, 24]. Micronization, a process that uses a supercritical fluid to change physicochemical aspects of substances, has been shown to improve the biological effects of substances such as N-acetylcysteine, resveratrol, and curcumin [5, 25–28]. In our previous studies, both curcumin and micronized curcumin (10 mg/kg, intraperitoneal) were unable to block the behavioral effects of acute restraint stress (90 minutes) [29] and unpredictable chronic stress (14 days) [28], while robust antioxidant effects were observed for micronized but not for its conventional preparation in adult zebrafish [28].

Larval stage zebrafish may occupy different ecological niches and face different environmental demands (e.g., type of predators, social behavior, preference for certain environments) when compared to adult zebrafish and therefore the behavioral phenotypes can be distinct [30]. In addition, zebrafish larvae have a more permeable blood-brain barrier (BBB) than adults and increased absorption of small molecules in water as well [31]. Thus, considering the pharmacokinetic limitations of curcumin, tests in larvae may respond differently from those observed in adults, and introducing the compound in water can better evidence the effects of micronization on the bioavailability. Therefore, this study aimed to compare the effects of exposure to conventional and micronized curcumin on locomotor, anxiety, cognitive, and biochemical parameters in zebrafish larvae.

## MATERIALS AND METHODS

### Drugs

Curcumin was obtained from Sigma-Aldrich™ (CAS 458-37-7) (St. Louis, MO, USA). Its micronization was carried out at the Laboratory of Thermodynamics and Supercritical Technology (LATESC) of the Departmento de Engenharia Química e de Alimentos (EQA) at Universidade Federal de Santa Catarina (UFSC), with the enhanced dispersion solution by supercritical fluids (SEDS) according to SACHETT et al. (2021a). Both curcumin preparations were dissolved in 0.1% dimethyl sulfoxide anhydrous (DMSO) obtained from Sigma-Aldrich (CAS 67-68-5) and diluted in the system water. Reagents used for biochemical assays were obtained from Sigma Aldrich (St. Louis, MO, USA): bovine serum albumin (CAS Number 9048-46-8), 5,5′-dithiobis (2-nitrobenzoic acid) (DTNB) (CAS Number 69-78-3), thiobarbituric acid (TBA) (CAS Number: 504-17-6), and trichloroacetic acid (TCA) (CAS Number: 76-03-9). Absolute ethanol (CAS Number: 64-17-5) was obtained from Merck KGaA (Darmstadt, Germany).

### Animals

All procedures were approved by the institutional animal welfare and ethical review committee at the Universidade Federal do Rio Grande do Sul (UFRGS) (approval #35279/2018). The experiments are reported in compliance with the ARRIVE guidelines 2.0 [32]. The embryos used were obtained from mating wild-type zebrafish (*Danio rerio*) from the colony of the Departamento de Bioquímica at UFRGS. The animals were maintained in recirculating systems (Zebtec™, Tecniplast, Italy) with reverse osmosis filtered water equilibrated to reach the species standard parameters including temperature (28 ± 2 °C), pH (7 ± 0.5), conductivity and ammonia, nitrite, nitrate, and chloride levels. The total organic carbon concentration was 0.33 mg/L and the total alkalinity (as carbonate ion) was 0.030 mEq/L. The animals were kept with a light/dark cycle of 14/10 h. The system water used in the experiments was obtained from a reverse osmosis apparatus (18 MOhm/cm) and was reconstituted with marine salt (Crystal SeaTM, Marinemix, Baltimore, MD, USA) at 0.4 ppt. For the breeding, in a breeding tank, females and males (1:2) were separated overnight by a transparent barrier, which was removed after the lights went on the following morning. The fertilized eggs retained in the bottom of the fitted tank were collected 1 hour after fertilization and the viable embryos were sanitized. They were randomly relocated in 24-well plates (4 embryos per well with 2 mL of system water), and kept in a biochemical oxygen demand incubator (BOD), on the same shelf to ensure the same lighting and housing pattern. The temperature was constant (28 °C) with a light/dark period of 14/10 hours until the 7^th^ day post-fertilization (dpf). At 7 dpf, the larvae were exposed to treatments and submitted to behavioral or biochemical tests. All experiments were performed in duplicate to confirm the results. Since zebrafish larvae can absorb all the necessary nutrients through the yolk sac up to 7 dpf, it was not necessary to feed the animals during the experiment. Embryos were evaluated daily to monitor mortality and hatching rates. At the end of the experiments, the zebrafish larvae were euthanized by immersion in cold water (0 to 4 °C) for 20 minutes until the cessation of the movements, to ensure death by hypoxia according to the AVMA Guidelines for the Euthanasia of Animals [33].

### Experimental design

The experiments were performed according to Figure 1. Two different experimental sets were carried out for the behavioral (open tank and light/dark (n = 16), and aversive stimulus tests (n = 4)) and another for the biochemical tests (levels of non-protein thiols and thiobarbituric acid reactive substances) (n = 3, pool = 15).

**Figure 1.**
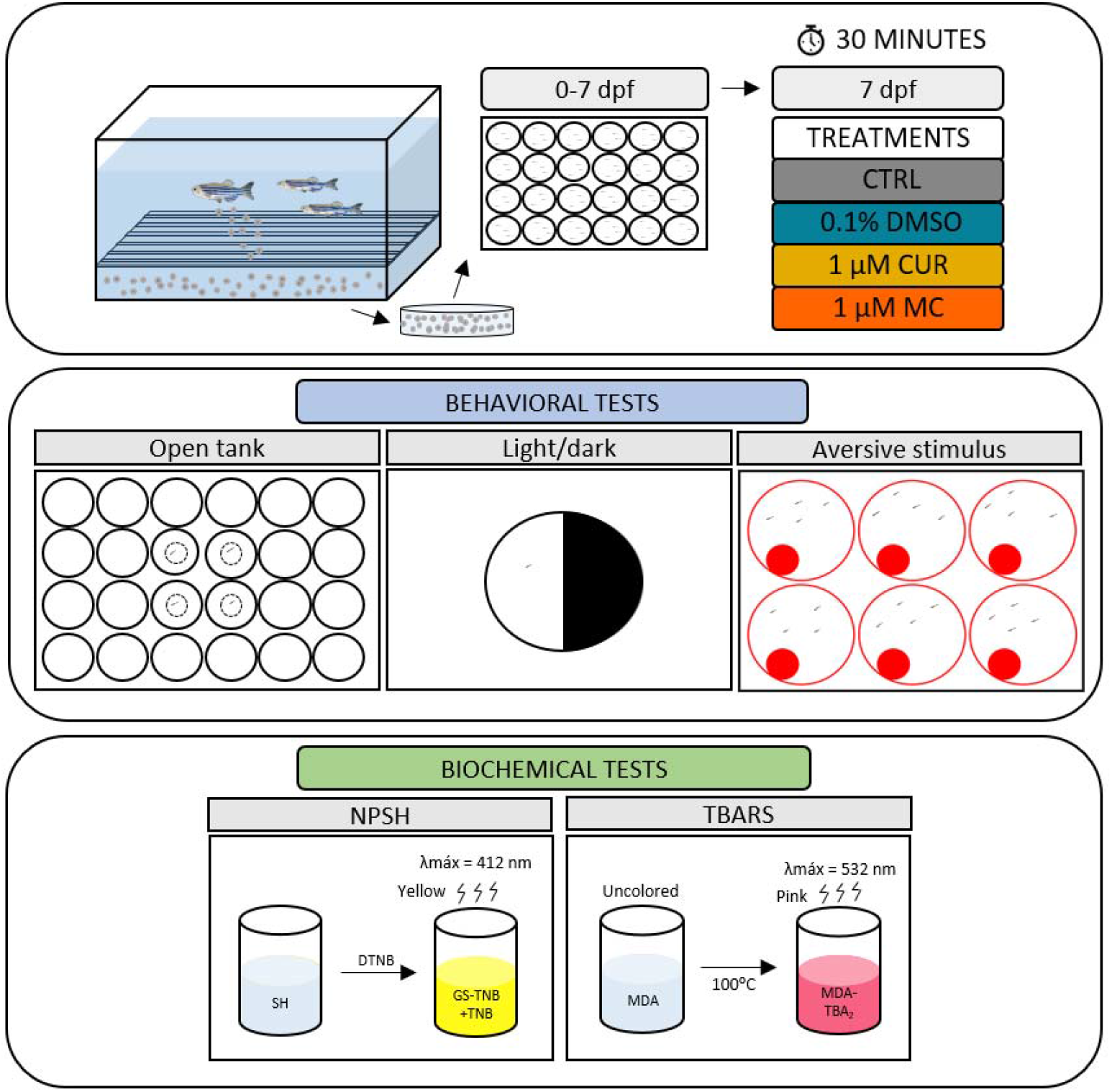
Experimental design. Two different sets of animals were used for the behavioral (set 1) and biochemical (set 2) tests. Initially, zebrafish larvae were randomly allocated to the experimental groups, 4 larvae per well, in a 24-well plate for the set 1 and 15 larvae per well, in a 6-well plate for set 2. At 7 dpf the larvae were exposed for 30 minutes to the following treatments: water (control), 0.1% DMSO (dilution vehicle), 1 μM CUR, or 1 μM MC. Subsequently, set 1 was tested in the open tank, light/dark (n = 16), and aversive stimulus tests (n = 4), and set 2 was assayed for NPSH and TBARS levels (n = 3, pool = 15). Dpf (days post-fertilization), CTRL (control), DMSO (dimethyl sulfoxide), CUR (curcumin), MC (micronized curcumin), NPSH (non-protein thiols) and TBARS (thiobarbituric acid reactive substances).

At 7 dpf, the animals were randomly allocated to the experimental groups, 4 larvae per well, in a 24-well plate (2 mL) for the behavioral tests and 15 larvae per well, in a 6-well plate (5 mL) for biochemical tests. The animals were allocated following block randomization procedures to counterbalance the housing plate (2 different plates). The larvae were exposed for 30 minutes to the following treatments: water (control), 0.1% DMSO (dilution vehicle), 1 μM curcumin (CUR), or 1 μM micronized curcumin (MC). Both curcumin preparations were diluted in 0.1% DMSO. All treatments were administered by immersion, being the substances dissolved in the system water. Concentrations were pre-established based on the literature [5]. All experimenters who analyzed the data were blind to the treatment groups.

### Open tank test (OTT)

To assess behavioral effects of treatments on locomotor and exploratory parameters the OTT was performed [2, 15, 34]. After 30 minutes of exposure to the treatments, the 7 dpf larvae were transferred individually to a 24-well plate filled with 2 mL of system water. The behavior was recorded from the top view and analyzed for 5 min, following 1 min of acclimation. The videos were analyzed with the ANYmaze® software (Stoelting Co., USA). The apparatus was virtually divided into two areas for video analysis: the central area of 8 mm in diameter and the periphery of 8 mm. The following parameters were quantified: total distance (m), crossings (transitions between the areas of the well), absolute turn angle (°), immobility time (s), and center time (time spent in the center area of the well) (s).

### Light/dark test (LDT)

After the OTT, the LDT was performed to assess anxiety parameters [30]. The larvae were individually placed on the light (white) side of the light/dark arena, which consists of a petri dish (60 × 15 mm) divided equally into two areas, light (white) and dark (black), filled with 15 mL of system water. The animals were recorded for 5 minutes, and the behavior was analyzed with BORIS® software. The time spent in the light area was measured.

### Aversive stimulus test (AST)

After the LDT, the avoidance capacity and cognitive responses to a visual stimulus were evaluated using an aversive task [35, 36]. The larvae were placed in 6-well plates (4 larvae per well, n = 4, filled with 5 mL of system water. The plates were placed on an LCD screen and exposed to an aversive visual stimulus. The video presented a red-filled circle that oscillates between the two ends of the well for 5 minutes. A habituation phase was carried out for 2 minutes before the stimulus started. The larvae were recorded and the time spent in the half without stimulation was evaluated. Animals that remained in the area with stimulation were considered to have cognitive and avoidance capacity dysfunction.

### Biochemical parameters

To assess the oxidative status, 30 minutes after exposure to treatments, biochemical tests were performed. Each pool of 15 larvae was simultaneously euthanized in a Petri dish with ice water at 0-4 °C. Afterward, all larvae were placed in a microtube (remaining water was removed) and 350 µL of phosphate-buffered saline (PBS, pH 7.4, Sigma-Aldrich) was added, then homogenized 60x with a pistil and centrifuged at 10,000 g at 4 °C in a refrigerated centrifuge. The supernatants were collected and kept in microtubes on ice until the assays were performed. The protein content was quantified according to the coomassie blue method using bovine serum albumin as a standard [37]. The detailed protocol for protein quantification is available at protocols.io [38].

### Non-protein thiols (NPSH)

The content of NPSH in the samples was determined by mixing equal volumes of the samples (50 µg of proteins) and TCA (6%). The mix was centrifuged (10,000 g, 10 min at 4 °C) and the supernatants were added to potassium phosphate buffer (TFK, 1 M) and DTNB (10 mM). After 1 h, the absorbance was measured at 412 nm. The detailed protocol is available at protocols.io [39].

### Thiobarbituric acid reactive substances (TBARS)

Lipid peroxidation was evaluated by quantifying the production of TBARS. Samples (50 µg of proteins) were mixed with TBA (0.5%) and TCA (20%). The mixture was heated at 100 °C for 30 min. The absorbance was measured at 532 nm in a microplate reader. MDA (2 mM) was used as the standard. The detailed protocol is available at protocols.io [40].

### Statistical analysis

We calculated the sample size to detect an effect size of 0.5 with a power of 0.9 and an alpha of 0.05 using G*Power 3.1.9.7 for Windows. The total distance traveled in OTT was defined as the primary outcome. The total sample size was 64, n = 16 animals per experimental group.

The normality and homogeneity of variances were confirmed for all data sets using D’Agostino-Pearson and Levene tests, respectively. Results were analyzed by one-way ANOVA followed by Tukey’s *post hoc* test when appropriate. The outliers were defined using the ROUT statistical test using the total distance traveled as the primary outcome and were removed from the analyses. This resulted in 5 outliers being removed from the OTT test (1 animal from CTRL, DMSO, and MC and 2 animals from CUR groups). The house plate effect was tested in all comparisons and no effect was observed, so the data were pooled.

Data are expressed as mean ± standard deviations of the mean (S.D.). The level of significance was set at p<0.05. Data were analyzed using IBM SPSS Statistics version 27 for Windows and the graphs were plotted using GraphPad Prism version 8.0.1 for Windows.

## RESULTS

### Behavioral parameters

The effects of CUR and MC on behavioral parameters in larvae zebrafish submitted to OTT are presented in Figure 2. Acute MC exposure increased the distance (F_3,55_ = 5.142, p=0.0033) (Fig. 2A) and absolute turn angle (F_3,55_ = 3.688, p=0.0378) (Fig. 2C). One-way ANOVA revealed a significant effect on the immobility time (Fig. 2D); however, no significant effects were observed in *post hoc* analysis. These results indicate an increase in locomotion and absolute turn angle in zebrafish larvae exposed to MC. There was no statistical difference of any intervention in the number of crossings and center time. Table 1 summarizes the one-way ANOVA analysis.

**Figure 2.**
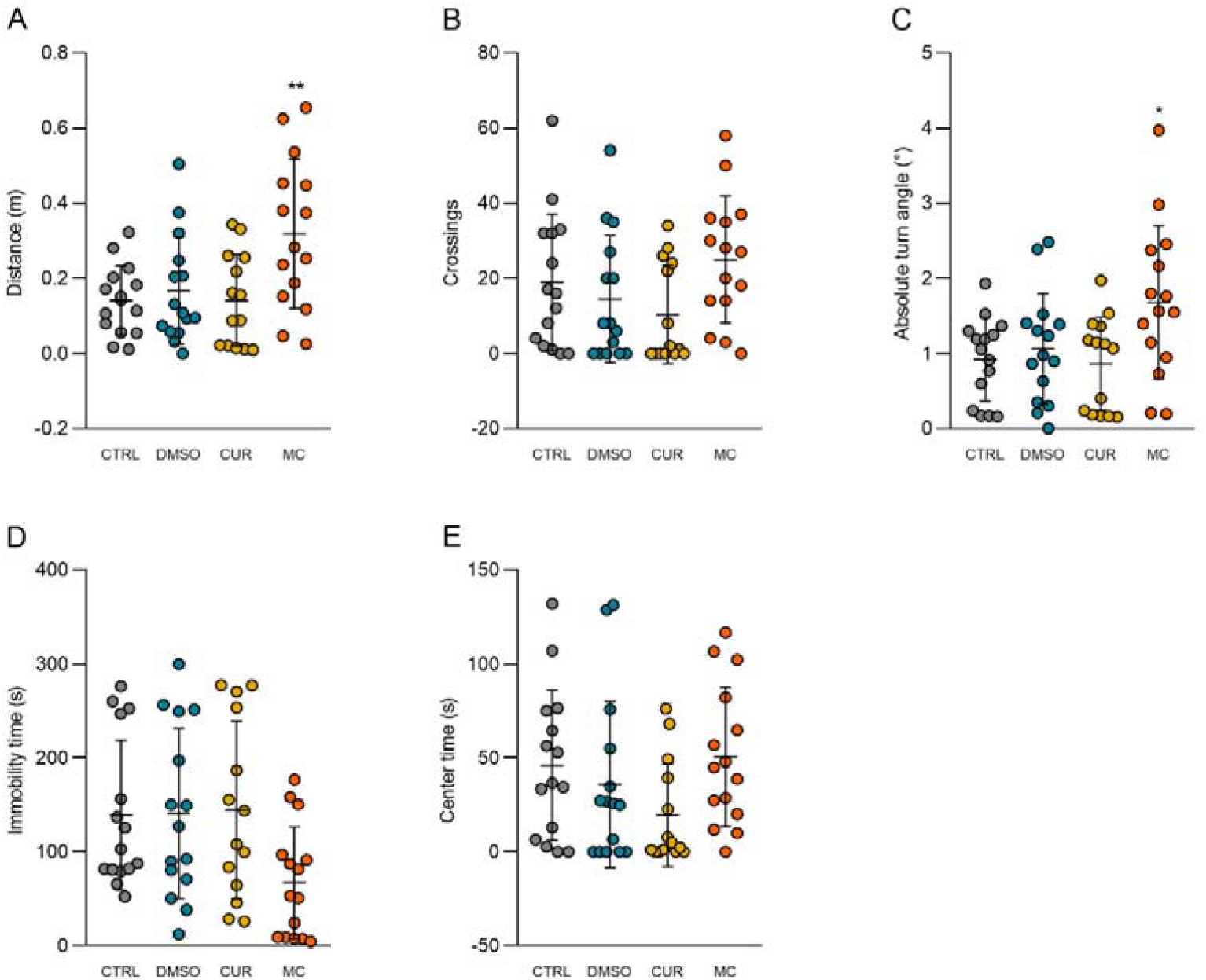
Effects of CUR and MC (1 µM) on behavioral parameters in the open tank test. (A) distance, (B) crossings, (C) absolute turn angle, (D) immobility time, and (E) center time. Data are expressed as mean ± S.D. One-way ANOVA/Tukey. n = 14-15. ^*^p<0.05 vs. CTRL. ^**^p<0.005 vs. CTRL. CTRL (control), DMSO (dimethyl sulfoxide), CUR (curcumin), MC (micronized curcumin).

**Table 1.** Summary of the one-way ANOVAs for the open tank, light/dark, aversive and biochemical tests.

| Test | Dependent variable | F-value | DF | P-value |
| --- | --- | --- | --- | --- |
| Open tank test | Distance | 5.142 | 3,55 | **<0.05** |
|  | Crossings | 2.109 | 3,55 | >0.05 |
|  | Absolute turn angle | 3.688 | 3,55 | **<0.05** |
|  | Immobility time | 3.079 | 3,55 | **<0.05** |
|  | Center time | 1.907 | 3,55 | >0.05 |
| Light/dark test | Time in the lit side | 0.9696 | 3,60 | >0.05 |
| Aversive stimulus test | % of animals in the non-stimulus area | 1.830 | 3,12 | >0.05 |
| Biochemical test | NPSH | 1.250 | 3,8 | >0.05 |
|  | TBARS | 0.5782 | 3,8 | >0.05 |

Significant effects ( $p < 0.05$ ) are given in bold font. DF (degrees of freedom), NPSH (non-protein thiols), TBARS (thiobarbituric acid reactive substances).

There was no statistical difference of any intervention in the time in the lit side in the LDT (Fig. 3), neither in percent of the animals in non-stimulus area in the AST (Fig. 4).

**Figure 3.**
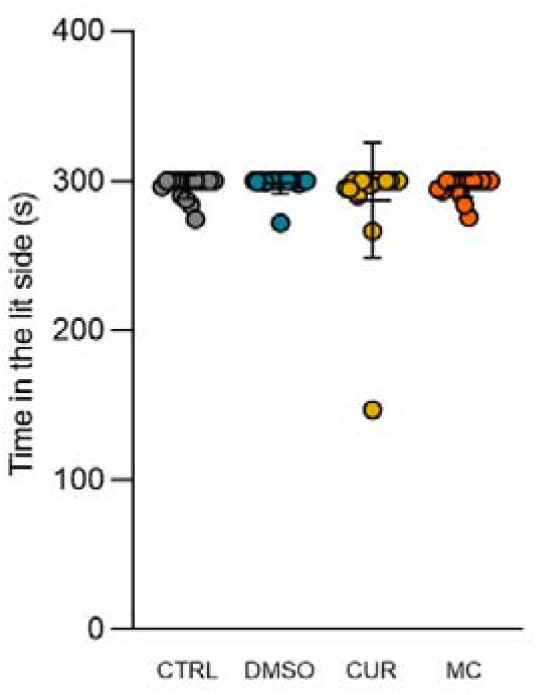
Effects of CUR and MC (1 µM) on behavioral parameters in the light/dark test. Data are expressed as mean ± S.D. One-way ANOVA. n = 16. CTRL (control), DMSO (dimethyl sulfoxide), CUR (curcumin), MC (micronized curcumin).

**Figure 4.**
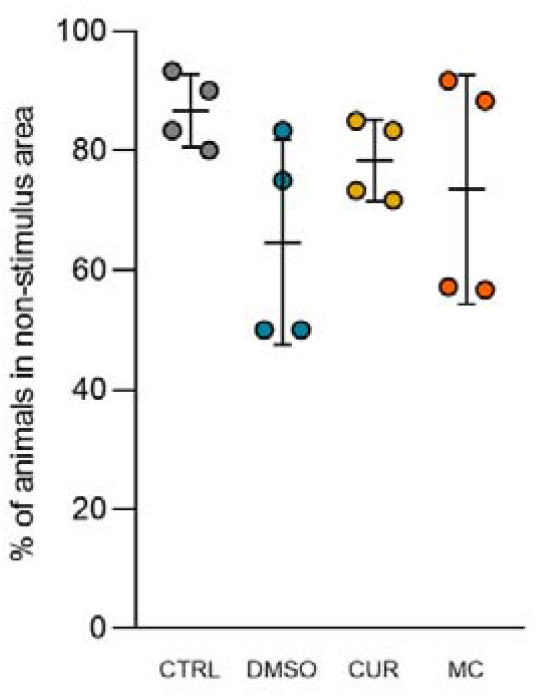
Effects of CUR and MC (1 µM) on behavioral parameters in the aversive test. Data are expressed as mean ± S.D. One-way ANOVA. n = 4. CTRL (control), DMSO (dimethyl sulfoxide), CUR (curcumin), MC (micronized curcumin).

### Biochemical parameters

The effects of CUR and MC on oxidative status parameters in zebrafish larvae are presented in Figure 5. There was no statistical difference of any intervention in the non-protein thiols and thiobarbituric acid reactive substances levels.

**Figure 5.**
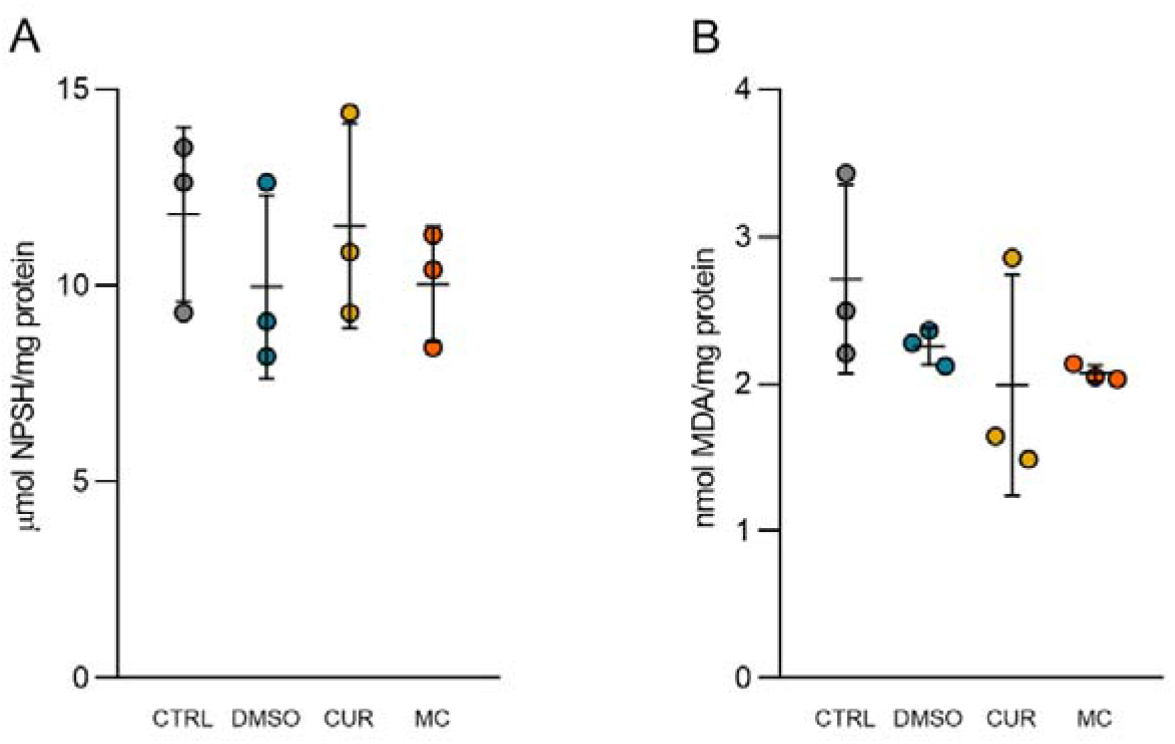
Effects of CUR and MC (1 µM) on biochemical parameters. (A) NPSH and (B) TBARS. Data are expressed as mean ± S.D. One-way ANOVA. n = 3. CTRL (control), DMSO (dimethyl sulfoxide), CUR (curcumin), MC (micronized curcumin), MDA (malondhyaldeide), NPSH (non-protein thiols), TBARS (thiobarbituric acid reactive substances).

## DISCUSSION

In this study, MC increased locomotion and absolute turn angle in the open tank test. However, there were no significant differences in the anxiety parameters in the light/dark or cognitive alterations in the aversive stimulus tests by any treatment. Also, there were no changes in the oxidative status of zebrafish larvae exposed to CUR or MC.

Essential for the survival of zebrafish, locomotion is a complex behavior controlled by the activity of various reticulospinal neurons of the brain stem along with descending vestibulospinal or neuromodulatory projections, whose activities depend on the integrity of brain function, nervous system development, and visual pathways [16, 41, 42]. The distance traveled and the pattern of movement of zebrafish larvae are relevant parameters to understand the neurobehavioral effects of the interventions [16]. The absolute turn angle is a measure of turning irrespective of its direction, which correlates with the motor pattern and is used to evaluate the motor coordination [43, 44]. In this study, MC increased the distance traveled and the absolute turn angle. Similar to our results, acute administration of neuroactive drugs that indirectly alter dopaminergic (DA) signaling as ethanol, d-amphetamine, and cocaine, increased locomotion in zebrafish larvae, and these effects were also observed in mammals [11, 12, 45]. Both the D_1_ and D_2_ selective agonists (SKF-38393 and quinpirole, respectively) increased locomotor activity, while the selective antagonists (SCH-23390 and haloperidol, D_1_- and D_2_-antagonists respectively) decreased locomotion [12]. Also, ethanol and quinpirole increased absolute turn angle while clonazepam, haloperidol, and valproic acid had the opposite effect [43, 44, 46, 47]. Although speculative, the effects of MC on the parameters above may be related to the modulation of dopaminergic transmission.

Some studies have suggested that the antidepressant effects of curcumin are associated with the modulation of dopamine pathways. In mice, acute curcumin (5 and 10 mg/Kg, p.o. and 20–80 mg/kg, i.p.) produced a significant inhibition of the immobility in the tail suspension test, forced swimming, and locomotor test. At the same time, it increased dopamine, serotonin, and noradrenaline levels, inhibiting both monoamine oxidase types in the mouse brain [48, 49]. In our study, MC affected locomotion, suggesting that the micronization process increases the psychostimulant actions of curcumin in zebrafish larvae.

In the light-dark, zebrafish larvae show white preference (black avoidance). In this test, diazepam increases the time spent in the black side [30]. Our results indicate that curcumin does not present anxiolytic effects in zebrafish larvae since there was no change in time spent in the light side in LDT [30] neither in time spent in the periphery area (thigmotaxis) in the OTT [15]. Moreover, curcumin does not change the percentage of animals in the non-stimulus area in the AST. Avoidance behavior in the 7-day-old larvae is important to detect a predator in the environment and may be also an indirect measure of anxiety and cognition [15]. Indeed, curcumin does not appear to have an acute anxiolytic effect or to impair avoidance behavior. In mice, curcumin (20 mg/kg) did not demonstrate any significant anxiolytic effects in the elevated plus-maze, open field test, and forced swim test, and no interaction of curcumin at the benzodiazepine site of the GABA_A_ receptor was observed [50]. In OTT, larvae show significantly less thigmotaxis behavior after exposure to the anxiolytic diazepam but not to the antidepressant fluoxetine [15]. In the same study, the diazepam-treated larvae showed an increased avoidance, unlike fluoxetine and caffeine-exposed larvae that displayed a decrease on this parameter. The differences between diazepam and fluoxetine are expected since these drugs target different neuronal signaling pathways [15]. Perhaps, for these same reasons, there were no observed effects of curcumin in these behavioral tests.

Although curcumin has antioxidant effects in several studies both *in vitro* and *in vivo* [28, 51, 52], we did not find an increase in antioxidant protection related to the non-enzymatic antioxidant GSH (represented by NPSH), nor a decrease in basal lipid peroxidation levels (represented by TBARS) in zebrafish larvae. In our previous study, acute exposure to curcumin and micronized curcumin had no significant antioxidant effects in adult zebrafish. We hypothesize that these effects are only observed when there is an increase in the production of reactive oxygen species or oxidative damage, in higher concentrations of the curcumin and/or for a longer period of exposure in adult and larval zebrafish [17, 28].

## CONCLUSION

In this study, MC increased locomotion and absolute turn angle but did not change anxiety, cognitive, and oxidative status outcomes in zebrafish larvae. We infer that the micronization process increases the stimulant effects of curcumin, which could be related to the indirect modulation of dopaminergic signaling. However, studies with dopaminergic antagonists and extending the exposure time and concentrations of the treatments should be performed to better characterize these results.

## Acknowledgments

We thank the Coordenação de Aperfeiçoamento de Pessoal de Nível Superior - Brasil (CAPES), Conselho Nacional de Desenvolvimento Científico e Tecnológico (CNPq, proc. 303343/2020-6), and Pró-Reitoria de Pesquisa (PROPESQ) at Universidade Federal do Rio Grande do Sul (UFRGS) for funding.

## Author contributions

All authors had full access to all the data in the study and take responsibility for the integrity of the data and the accuracy of the data analysis. Conceptualization, A.S., and A.P.; Methodology, A.S., R.B., C.G.R., M.G-L., L.M.B., G.P.S.A., A.P.H. J.V.O., A.M.S., A.P.; Investigation, A.S., R.B., C.G.R., M.G-L., L.M.B., G.P.S.A.; Formal Analysis, R.B., C.G.R., A.P. and A.P.H.; Resources, J.V.O., A.M.S., A.P.; Writing - Original Draft, A.S.; Writing - Review & Editing, A.S., R.B., C.G.R., M.G-L., L.M.B., G.P.S.A., A.P.H., J.V.O., A.M.S., A.P.; Supervision, A.P.; Funding Acquisition, A.P.

## Data availability

The datasets generated in the current study are available from the corresponding author on reasonable request.

## REFERENCES

1. MacRae CA, Peterson RT (2015) Zebrafish as tools for drug discovery. Nat Rev Drug Discov 14:721–731. https://doi.org/10.1038/nrd4627

2. Benvenutti R, Marcon M, Reis CG, et al (2018) N-acetylcysteine protects against motor, optomotor and morphological deficits induced by 6-OHDA in zebrafish larvae. PeerJ 6:e4957. https://doi.org/10.7717/peerj.4957

3. Benvenutti R, Gallas-Lopes M, Sachett A, et al (2021) How do zebrafish (Danio rerio) respond to MK-801 and amphetamine? Relevance for assessing schizophrenia-related endophenotypes in alternative model organisms. J Neurosci Res n/a: https://doi.org/10.1002/jnr.24948

4. Benvenutti R, Gallas-Lopes M, Marcon M, et al (2021) Glutamate Nmda Receptor Antagonists With Relevance To Schizophrenia: A Review Of Zebrafish Behavioral Studies. Curr Neuropharmacol. https://doi.org/10.2174/1570159X19666210215121428

5. Bertoncello KT, Aguiar GPS, Oliveira JV, Siebel AM (2018) Micronization potentiates curcumin’s anti-seizure effect and brings an important advance in epilepsy treatment. Sci Rep 8:2645. https://doi.org/10.1038/s41598-018-20897-x

6. Lee J, Freeman J (2016) Embryonic exposure to 10 μg L(-1) lead results in female-specific expression changes in genes associated with nervous system development and function and Alzheimer’s disease in aged adult zebrafish brain. Met Integr Biometal Sci. https://doi.org/10.1039/c5mt00267b

7. Mocelin R, Marcon M, da Rosa Araujo AS, et al (2019) Withdrawal effects following repeated ethanol exposure are prevented by N-acetylcysteine in zebrafish. Prog Neuropsychopharmacol Biol Psychiatry 93:161–170. https://doi.org/10.1016/j.pnpbp.2019.03.014

8. Patton EE, Zon LI, Langenau DM (2021) Zebrafish disease models in drug discovery: from preclinical modelling to clinical trials. Nat Rev Drug Discov 20:611–628. https://doi.org/10.1038/s41573-021-00210-8

9. Piato ÂL, Capiotti KM, Tamborski AR, et al (2011) Unpredictable chronic stress model in zebrafish (Danio rerio): Behavioral and physiological responses. Prog Neuropsychopharmacol Biol Psychiatry 35:561–567. https://doi.org/10.1016/j.pnpbp.2010.12.018

10. Akhtar MT, Ali S, Rashidi H, et al (2013) Developmental effects of cannabinoids on zebrafish larvae. Zebrafish 10:283–293. https://doi.org/10.1089/zeb.2012.0785

11. Irons TD, MacPhail RC, Hunter DL, Padilla S (2010) Acute neuroactive drug exposures alter locomotor activity in larval zebrafish. Neurotoxicol Teratol 32:84–90. https://doi.org/10.1016/j.ntt.2009.04.066

12. Irons TD, Kelly PE, Hunter DL, et al (2013) Acute administration of dopaminergic drugs has differential effects on locomotion in larval zebrafish. Pharmacol Biochem Behav 103:792–813. https://doi.org/10.1016/j.pbb.2012.12.010

13. Parker B, Connaughton VP (2007) Effects of nicotine on growth and development in larval zebrafish. Zebrafish 4:59–68. https://doi.org/10.1089/zeb.2006.9994

14. Raftery TD, Isales GM, Yozzo KL, Volz DC (2014) High-Content Screening Assay for Identification of Chemicals Impacting Spontaneous Activity in Zebrafish Embryos. Environ Sci Technol 48:804–810. https://doi.org/10.1021/es404322p

15. Richendrfer H, Pelkowski SD, Colwill RM, Creton R (2012) On the edge: Pharmacological evidence for anxiety-related behavior in zebrafish larvae. Behav Brain Res 228:99–106. https://doi.org/10.1016/j.bbr.2011.11.041

16. Basnet RM, Zizioli D, Taweedet S, et al (2019) Zebrafish Larvae as a Behavioral Model in Neuropharmacology. Biomedicines 7:23. https://doi.org/10.3390/biomedicines7010023

17. Arteaga C, Boix N, Teixido E, et al (2021) The Zebrafish Embryo as a Model to Test Protective Effects of Food Antioxidant Compounds. Molecules 26:5786. https://doi.org/10.3390/molecules26195786

18. Endo Y, Muraki K, Fuse Y, Kobayashi M (2020) Evaluation of Antioxidant Activity of Spice-Derived Phytochemicals Using Zebrafish. Int J Mol Sci 21:E1109. https://doi.org/10.3390/ijms21031109

19. Kim S, Kim M, Kang M-C, et al (2021) Antioxidant Effects of Turmeric Leaf Extract against Hydrogen Peroxide-Induced Oxidative Stress In Vitro in Vero Cells and In Vivo in Zebrafish. Antioxid Basel Switz 10:112. https://doi.org/10.3390/antiox10010112

20. Zhang R, Zhao T, Zheng B, et al (2021) Curcumin Derivative Cur20 Attenuated Cerebral Ischemic Injury by Antioxidant Effect and HIF-1α/VEGF/TFEB-Activated Angiogenesis. Front Pharmacol 12:648107. https://doi.org/10.3389/fphar.2021.648107

21. Choo BKM, Kundap UP, Faudzi SMM, et al (2021) Identification of curcumin analogues with anti-seizure potential in vivo using chemical and genetic zebrafish larva seizure models. Biomed Pharmacother Biomedecine Pharmacother 142:112035. https://doi.org/10.1016/j.biopha.2021.112035

22. Muthuraman A, Thilagavathi L, Jabeen S, et al (2019) Curcumin prevents cigarette smoke extract induced cognitive impairment. Front Biosci Elite Ed 11:109–120. https://doi.org/10.2741/E850

23. Anand P, Kunnumakkara AB, Newman RA, Aggarwal BB (2007) Bioavailability of Curcumin: Problems and Promises. Mol Pharm 4:807–818. https://doi.org/10.1021/mp700113r

24. Yang K-Y, Lin L-C, Tseng T-Y, et al (2007) Oral bioavailability of curcumin in rat and the herbal analysis from Curcuma longa by LC–MS/MS. J Chromatogr B 853:183–189. https://doi.org/10.1016/j.jchromb.2007.03.010

25. Aguiar GPS, Marcon M, Mocelin R, et al (2017) Micronization of N-acetylcysteine by supercritical fluid: Evaluation of in vitro and in vivo biological activity. J Supercrit Fluids 130:282–291. https://doi.org/10.1016/j.supflu.2017.06.010

26. Almeida ER, Lima-Rezende CA, Schneider SE, et al (2021) Micronized Resveratrol Shows Anticonvulsant Properties in Pentylenetetrazole-Induced Seizure Model in Adult Zebrafish. Neurochem Res 46:241–251. https://doi.org/10.1007/s11064-020-03158-0

27. Decui L, Garbinato CLL, Schneider SE, et al (2020) Micronized resveratrol shows promising effects in a seizure model in zebrafish and signalizes an important advance in epilepsy treatment. Epilepsy Res 159:106243. https://doi.org/10.1016/j.eplepsyres.2019.106243

28. Sachett A, Gallas-Lopes M, Benvenutti R, et al (2021) Curcumin micronization by supercritical fluid: in vitro and in vivo biological relevance. bioRxiv 2021.07.08.451641. https://doi.org/10.1101/2021.07.08.451641

29. Sachett A, Gallas-Lopes M, Benvenutti R, et al (2021) Non-micronized and micronized curcumin do not prevent the behavioral and neurochemical effects induced by acute stress in zebrafish. https://doi.org/10.1101/2021.10.11.463974

30. Steenbergen PJ, Richardson MK, Champagne DL (2011) Patterns of avoidance behaviours in the light/dark preference test in young juvenile zebrafish: A pharmacological study. Behav Brain Res 222:15–25. https://doi.org/10.1016/j.bbr.2011.03.025

31. Fleming A, Diekmann H, Goldsmith P (2013) Functional characterisation of the maturation of the blood-brain barrier in larval zebrafish. PloS One 8:e77548. https://doi.org/10.1371/journal.pone.0077548

32. Sert NP du, Hurst V, Ahluwalia A, et al (2020) The ARRIVE guidelines 2.0: Updated guidelines for reporting animal research. Br J Pharmacol 177:3617–3624. https://doi.org/10.1111/bph.15193

33. Leary S, Pharmaceuticals F, Underwood W, et al (2020) AVMA Guidelines for the Euthanasia of Animals: 2020 Edition. 121

34. Creton R (2009) Automated analysis of behavior in zebrafish larvae. In: Behav. Brain Res. https://pubmed.ncbi.nlm.nih.gov/19409932/. Accessed 1 Dec 2020

35. Nery LR, Silva NE, Fonseca R, Vianna MRM (2017) Presenilin-1 Targeted Morpholino Induces Cognitive Deficits, Increased Brain Aβ1-42 and Decreased Synaptic Marker PSD-95 in Zebrafish Larvae. Neurochem Res 42:2959–2967. https://doi.org/10.1007/s11064-017-2327-4

36. Pelkowski SD, Kapoor M, Richendrfer HA, et al (2011) A novel high-throughput imaging system for automated analyses of avoidance behavior in zebrafish larvae. Behav Brain Res 223:135–144. https://doi.org/10.1016/j.bbr.2011.04.033

37. Bradford MM (1976) A rapid and sensitive method for the quantitation of microgram quantities of protein utilizing the principle of protein-dye binding. Anal Biochem 72:248–254. https://doi.org/10.1006/abio.1976.9999

38. Sachett A (2020) Protein quantification protocol optimized for zebrafish brain tissue (Bradford method). https://doi.org/10.17504/protocols.io.bjnfkmbn

39. Sachett A, Gallas-Lopes M, Conterato GMM, et al (2021) Quantification of nonprotein sulfhydryl groups (NPSH) optimized for zebrafish brain tissue. In: protocols.io. https://www.protocols.io/view/quantification-of-nonprotein-sulfhydryl-groups-nps-bx8tprwn. Accessed 11 Oct 2021

40. Sachett A (2020) Quantification of thiobarbituric acid reactive species (TBARS) optimized for zebrafish brain tissue. https://doi.org/10.17504/protocols.io.bjp8kmrw

41. Brustein E, Brustein S-A, Buss R, et al (2003) Steps during the development of the zebrafish locomotor network. In: J. Physiol. Paris. https://pubmed.ncbi.nlm.nih.gov/14706693/. Accessed 1 Dec 2020

42. Drapeau P, Saint-Amant L, Buss RR, et al (2002) Development of the locomotor network in zebrafish. Prog Neurobiol 68:85–111. https://doi.org/10.1016/s0301-0082(02)00075-8

43. Blazina AR, Vianna MR, Lara DR (2013) The spinning task: a new protocol to easily assess motor coordination and resistance in zebrafish. Zebrafish 10:480–485. https://doi.org/10.1089/zeb.2012.0860

44. Tran S, Chow H, Tsang B, et al (2017) Zebrafish Are Able to Detect Ethanol in Their Environment. Zebrafish 14:126–132. https://doi.org/10.1089/zeb.2016.1372

45. Liu X, Lin J, Zhang Y, et al (2016) Effects of diphenylhydantoin on locomotion and thigmotaxis of larval zebrafish. Neurotoxicol Teratol 53:41–47. https://doi.org/10.1016/j.ntt.2015.11.008

46. Nabinger DD, Altenhofen S, Peixoto JV, et al (2021) Long-lasting behavioral effects of quinpirole exposure on zebrafish. Neurotoxicol Teratol 88:107034. https://doi.org/10.1016/j.ntt.2021.107034

47. Tran S, Facciol A, Gerlai R (2016) Alcohol-induced behavioral changes in zebrafish: The role of dopamine D2-like receptors. Psychopharmacology (Berl) 233:2119–2128. https://doi.org/10.1007/s00213-016-4264-3

48. Kulkarni SK, Bhutani MK, Bishnoi M (2008) Antidepressant activity of curcumin: involvement of serotonin and dopamine system. Psychopharmacology (Berl) 201:435. https://doi.org/10.1007/s00213-008-1300-y

49. Xu Y, Ku B-S, Yao H-Y, et al (2005) The effects of curcumin on depressive-like behaviors in mice. Eur J Pharmacol 518:40–46. https://doi.org/10.1016/j.ejphar.2005.06.002

50. Ceremuga TE, Helmrick K, Kufahl Z, et al (2017) Investigation of the Anxiolytic and Antidepressant Effects of Curcumin, a Compound From Turmeric (Curcuma longa), in the Adult Male Sprague-Dawley Rat. Holist Nurs Pract 31:193–203. https://doi.org/10.1097/HNP.0000000000000208

51. Ak T, Gülçin I (2008) Antioxidant and radical scavenging properties of curcumin. Chem Biol Interact 174:27–37. https://doi.org/10.1016/j.cbi.2008.05.003

52. Menon VP, Sudheer AR (2007) ANTIOXIDANT AND ANTI-INFLAMMATORY PROPERTIES OF CURCUMIN. In: Aggarwal BB, Surh Y-J, Shishodia S (eds) The Molecular Targets and Therapeutic Uses of Curcumin in Health and Disease. Springer US, Boston, MA, pp 105–125

